# The Tangent copy-number inference pipeline for cancer genome analyses

**DOI:** 10.1101/566505

**Authors:** Barbara Tabak, Gordon Saksena, Coyin Oh, Galen F. Gao, Barbara Hill Meyers, Michael Reich, Steven E. Schumacher, Lindsay C. Westlake, Ashton C. Berger, Scott L.b Carter, Andrew D. Cherniack, Matthew Meyerson, Rameen Beroukhim, Gad Getz

## Abstract

**Motivation:** Somatic copy-number alterations (SCNAs) play an important role in cancer development. Systematic noise in sequencing and array data present a significant challenge to the inference of SCNAs for cancer genome analyses. As part of The Cancer Genome Atlas (TCGA), the Broad Institute Genome Characterization Center developed the Tangent copy-number inference pipeline to generate copy-number profiles using single-nucleotide polymorphism (SNP) array and whole-exome sequencing (WES) data from over 10,000 pairs of tumors and matched normal samples. Here, we describe the Tangent pipeline, which begins with DNA sequencing data in the form of .bam files or raw SNP array probe-level intensity data, and ends with segmented copy-number calls to facilitate the identification of novel genes potentially targeted by SCNAs. We also describe a modification of Tangent, Pseudo-Tangent, which enables denoising through comparisons between tumor profiles when few normal samples are available.

**Results:** Tangent Normalization offers substantial signal-to-noise ratio (SNR) improvements compared to conventional normalization methods in both SNP array and WES analyses. The improvement in SNRs is achieved primarily through noise reduction with minimal effect on signal. Pseudo-Tangent also reduces noise when few normal samples are available. Tangent and Pseudo-Tangent are broadly applicable and enable more accurate inference of SCNAs from DNA sequencing and array data.

**Availability and Implementation:** Tangent is available at https://github.com/coyin/tangent and as a Docker image (https://hub.docker.com/r/coyin/tangent). Tangent is also the normalization method for the Copy Number pipeline in Genome Analysis Toolkit 4 (GATK4).

## 1 Introduction

Cancer often arises from the accumulation of somatic alterations in the genome, including point mutations, structural rearrangements, and copy number alterations (Weir *et al.*, 2004). Somatic copy number alterations (SCNAs) can have significant impact in activating oncogenes or inactivating tumor suppressor genes to drive the development of cancer (Beroukhim *et al.*, 2010; Zack *et al.*, 2013). In 2006, the NCI and NHGRI launched The Cancer Genome Atlas (TCGA) project to comprehensively characterize the genomic and molecular features of different cancer types (The Cancer Genome Atlas Research Network, 2013). TCGA collected samples from more than 11,000 cancer patients across 33 tumor types. The use of next-generation sequencing (NGS) and high-resolution microarrays allowed us to finely characterize SCNAs in cancer genomes and facilitate the discovery of novel genes that drive cancer (The Cancer Genome Atlas Network et al., 2013; Zack et al., 2013; Korn et al., 2008).

Standard approaches to detect somatic copy-number profiles involve determining DNA content at various sites across the genome in tumor samples, and comparing to normal samples. For example, array CGH or single nucleotide polymorphism (SNP) arrays are composed of DNA probes that match various genomic loci; signal intensities read from these arrays scale with sample DNA content at each locus (LaFramboise, 2009). Similarly, high-throughput sequencing enables determination of coverage levels at loci across the genome, also reflecting sample DNA content (Yoon *et al.*, 2009). Detection of somatic copy-number alterations (SCNAs) typically relies on determining the ratios between DNA content in tumor vs. normal samples across these loci, which aims to normalize the different affinities (either of probes or sequencing) associated with each locus.

Such analyses can be confounded by at least three sources of noise. First, stochastic variations result in random deviations between measurements of DNA content and true DNA content. This can be overcome by averaging measurements across adjacent loci (e.g. using segmentation algorithms; Venkatraman and Olshen, 2007) or by sequencing to greater average depth. Second, germline copy-number variations (CNVs) can be misinterpreted as SCNAs. This can be overcome by comparing tumor DNA to normal DNA from the same patient, or by masking common CNVs. Third, systematic errors can result from subtle differences between the experimental conditions that applied when generating sequencing or microarray data from tumors and their normal controls, which can affect the locus-specific affinities.

Despite rapid advancement in sequencing technologies and improvements in copy number tools that attempt to combat systematic noise, such as Control-FREEC, ExomeCNV, VarScan2, and CNVkit (Boeva *et al.*, 2012; Sathirapongsasuti *et al.*, 2011; Koboldt *et al.*, 2017; Talevich *et al.*, 2016; Rieber *et al.*, 2017; Zare *et al.*, 2017), filtering out systematic noise present in NGS and microarray data remains a significant challenge. Many of these tools use similar approaches to reduce systematic noise, either with matched case-control samples or with GC correction (Zhao *et al.*, 2013). While matched normal samples can sometimes approximate their tumors’ noise profiles, they are not always available, and during the sequencing process, many of them may be processed under conditions different from those of their corresponding tumors and therefore may not have similar noise profiles. And while GC-content bias constitutes a large component of systematic noise, GC correction does not target all sources of noise present in copy number data. Other potential sources of systematic noise include mappability biases across the genome and variability in experimental conditions during PCR amplification, cross-hybridization, or sample and library preparation. Thus, currently available tools do not adequately address these gaps in copy number analysis.

Here, we present Tangent, a copy number inference pipeline that aims to address these gaps by constructing noise profiles using a subset of normal samples to target all potential sources of systematic noise. The normal samples used for Tangent will ideally have been processed using the same experimental conditions as the tumor samples, but do not have to be from the same patients as the tumors. Our pipeline begins with either a whole-exome sequencing (WES) BAM file or raw probe-level intensity data and concludes with segmented copy number calls, processing data with special attention to noise reduction, artifact removal, and quality control. The Tangent pipeline can be applied to both WES and Affymetrix SNP Array 6.0, both of which have been the basis for data analyses in TCGA. Tangent can also be extended to other sequencing platforms. Additionally, we describe Pseudo-Tangent, an approach that uses signal-subtracted tumor data to augment standard normal data in the Tangent pipeline. Pseudo-Tangent is particularly useful when there is a limited number of normal samples that can be used for denoising. Tangent is the basis for copy-number normalization in the GATK4 CNV workflow available within Genome Analysis Toolkit 4 (GATK4; McKenna et al., 2010) and is available through Github and Docker.

## 2 Materials and Methods

### 2.1 Generation of raw coverage data

As input to Tangent, we generated raw coverage data from either Affymetrix SNP arrays or from WES. For SNP arrays, the procedure to generate raw coverage data is described in Supplementary Methods. For WES data, we used the GATK DepthOfCoverage tool on input .bam files to assess coverage from the input .bam file (Depristo *et al.*, 2011). DepthOfCoverage outputs values for a set of genomic loci (“intervals”) representing the hybrid capture targets. Interval files are available in Firecloud from the broad-firecloud-tutorials/Broad_MutationCalling_QC_Workflow_BestPractice workspace. Flow charts for each type of input data are presented in Supplementary Figure 1A-B.

**Figure 1.**
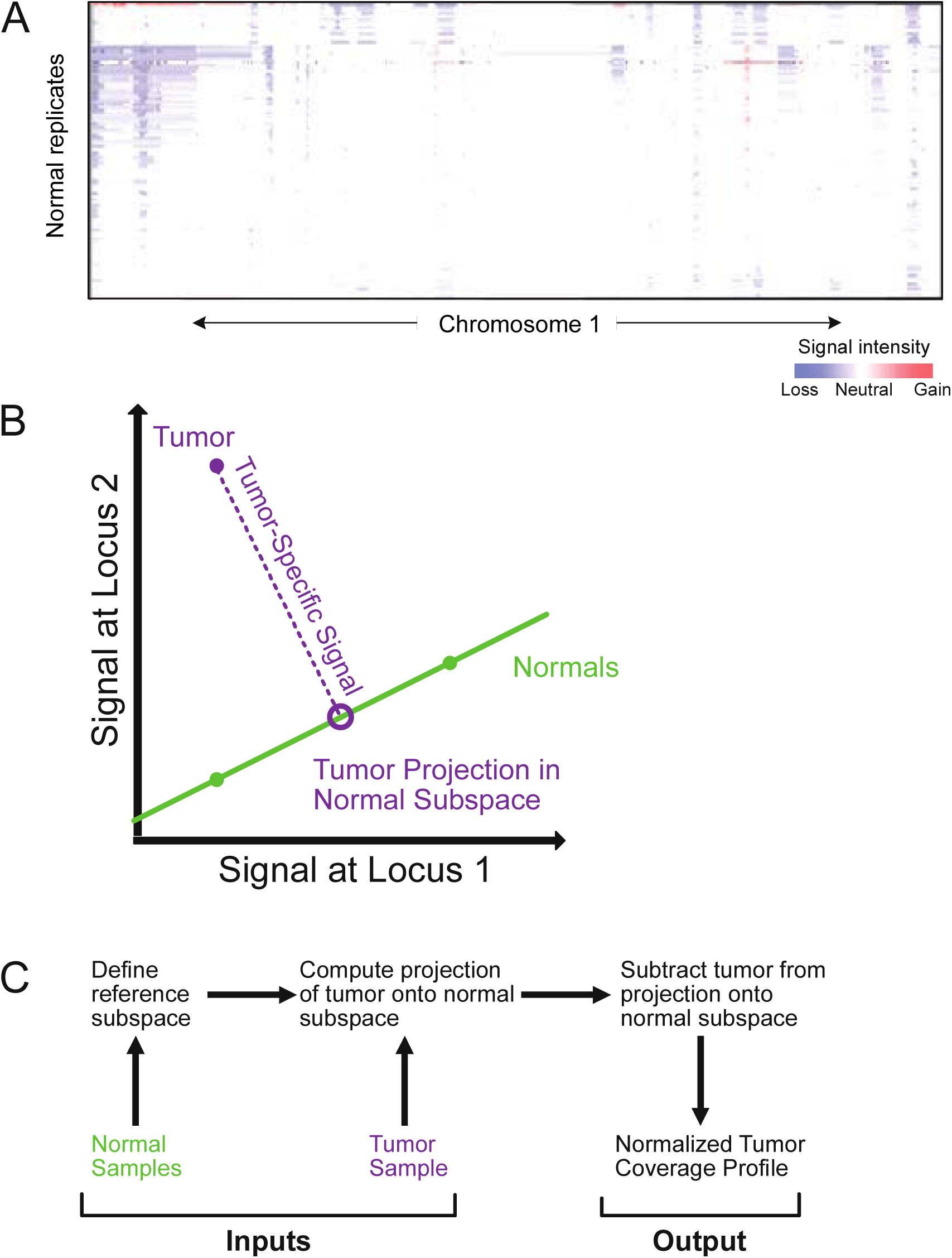
Overview of problem and method. (a) Segmented pre-normalized log_2_ copy-number ratios (low and high ratios indicated by blue and red, respectively) on replicates of DNA from the HCC1143BL immortalized lymphocyte (non-cancer) line across 110 batches in chromosome 1. As these variations are observed in the same DNA, they represent experimental artifacts. (b) Reduced, 2-dimensional representation of the Tangent methodology. For each tumor (purple) we compute its projection onto a lower-dimensional subspace defined by normal samples (green) profiled in parallel with the tumors. Signal representing somatic copy-number alterations is contained within the difference between the tumor and its projection. (c) Flowchart describing the steps of Tangent normalization.

### 2.2 Tangent Normalization

Tangent assumes that systematic noise, after log-transformation, is distributed according to a similar additive pattern in tumor samples as in normal samples. (We use log_2_ copy-ratios because we have found that this representation works well for noise reduction [data not shown], suggesting that much of the observed noise is multiplicative.) Therefore, to minimize systematic noise, we can subtract estimated noise profiles individually calculated for each tumor using data from normal samples. Specifically, we compute the projection of each tumor in a lower-dimensional subspace spanned by the coverage profiles of the normal samples and then subtract that projection from the raw log_2_-transformed copy number profile of the tumor. This difference is the Tangent-normalized coverage profile for that tumor.

For *i* ∈ {1,2,3, … *n*_*N*_ } where *n_N_* is the number of normal samples, the i^th^ normal sample is represented as a vector, *N*_*i*_, of log2 copy-ratio intensities in genomic order, with each coordinate corresponding to one of the non-CNV probes. The noise space, ***N***, is defined as the (*n*_*N*_ – 1)-dimensional plane containing the vectors 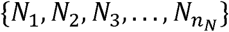 . Note that *n*_*N*_ – 1 << *M*, where *M* equals the dimension of the ambient (log2 copy-ratio) coordinate space or equivalently, the number of markers not excluded as poor quality or potential CNVs. Similarly, for *j* ∈ {1,2,3,…, *n*_*T*_ }, where *n*_*T*_ is the number of tumor samples, *T*_*j*_ represents the j^th^ tumor sample in the same format as *N*_*i*_. A constructed normal profile that closely matches the noise profile for a tumor *T*_*j*_ is determined as the point in ***N*** that is closest to *T*_*j*_ using a Euclidean metric, i.e. the projection, *p*(*T*_*j*_), of *T*_*j*_ on ***N***. The resulting normalization of *T*_*j*_ is set to the residual, *T*_*j*_ – *p*(*T*_*j*_).

The projection *p*(*T*_*j*_) can be computed directly using standard linear algebra techniques. A rigid transformation of Euclidean marker space prior to normalization does not alter the resulting normalization of *T*_*j*_. In particular, an appropriate translation of the Euclidean space ensures that ***N*** passes through the origin and forms a vector subspace of Euclidean space, in which the normal vectors now reflect the deviation from the typical normal (i.e. the noise). After projection to ***N***, the noise profile for each sample can be expressed as a linear combination of *n*_*N*_-1 translated normal vectors. This noise profile, that is closest to the tumor, is then subtracted from the tumor signal to obtain a “clean” signal.

We include both male and female normal samples, which differ in the number of copies of X. The inclusion of the X chromosome in Tangent Normalization requires special treatment to ensure that the distance from a tumor to a normal reflects noise differences, without being artificially inflated due to gender difference. Additionally, we must take into account that the normalization, *T*_*j*_ – *p*(*T*_*j*_), of *T*_*j*_ could potentially alter the apparent chromosomal copy number of X, due to the fact that *p*(*T*_*j*_) is a weighted average of copy ratios from both male and female samples. To address these issues, we include in our reference plane a theoretical normal with copy-number precisely two throughout the autosomes and one throughout the X chromosome. Tangent normalization against this expanded collection of normal samples will adjust the copy-profile of X for any sample, regardless of gender, to a mean level with ∼2 copies of X. The ensuing analysis can detect focal SCNAs within X, but discounts whole-chromosome changes of X. Currently, the Y chromosome is excluded from Tangent Normalization. Use of gender-matched normals may enable recovery of whole-chromosome SCNAs involving X.

The large number of reference normal samples presents computational challenges as the projection matrix depends on the computation of the pseudo-inverse of an *M* x *n*_*N*_ matrix (∼1.5e6 x 3000). To address this issue, we mimic Gram-Schmidt orthogonalization, but on a blockwise level, and decompose the reference plane into orthogonal blocks so that the projection, *p*(*T*_*j*_), can be computed on a block-by-block basis with only one block in memory at a time. Each block of data represents approximately 250 normal samples, typically from multiple batches. The orthogonalization process replaces the *i*^*th*^ block of normal data by its Tangent Normalization against blocks 1 through *i* – 1. When a new batch is processed, an additional block is added using the normal samples from the batch at hand, which are themselves first normalized against the reference normal samples.

### 2.3 Pseudo-Tangent

In the first step of Pseudo-Tangent, we use Tangent with a small set of normals to define the reference subspace and Circular Binary Segmentation (CBS) (Venkatraman and Olshen, 2007) to generate a tentative copy number profile for each tumor. In the second step, we subtract these tentative profiles from their original log-transformed tumor profiles in order to generate a corresponding pseudo-normal profile for each tumor input (keeping only deviations from the CBS segment values). In the penultimate step, the tumors are partitioned into *n* approximately equal subsets, and then each subset is Tangent-normalized against a reference subspace of pseudo-normals generated from tumors in that subset’s complement. The partition parameter *n* is a user-controlled parameter that is inversely related to the cardinality of each subset. Finally, CBS is used to convert the resulting Pseudo-Tangent-normalized coverage profiles into segmented copy-number calls in the form of log2 copy-ratios (Supplementary Figure S1C).

An optional step we take in Pseudo-Tangent is to perform truncated singular value decomposition (tSVD) on the entire collection of pseudo-normal profiles before partitioning the tumors and normalizing their coverages against the pseudo-normal profiles. This step limits the dimensionality of our pseudo-normal reference subspace and constrains us to a smaller number of eigenvectors to describe our pseudo-normal noise distribution.

### 2.4 Comparisons against other normalization methods

When comparing Tangent normalization to other normalization methods, we opted to exclude the X and Y chromosomes from our analyses so that differences in their handling of the sex chromosomes would not affect their performances. For similar reasons, we excluded CNV probes that map to known germline copy-number polymorphisms or other regions where, due to errors in the experimental platform, data across normals vary widely (Supplementary Table 1). To normalize using matched normals, we subtracted the log_2_ ratios of each matched normal from its corresponding tumor. For tumors with more than one matched normal (blood or normal tissue sample), the matched blood sample was preferred over the matched normal tissue sample. To normalize using the five nearest normals approach, we subtracted from each tumor the mean of the five normals closest to it based on Euclidean distance (Beroukhim *et al.*, 2007).

To normalize using the average normal method with WES data, we first averaged the coverage at each interval across the entire panel of normal samples to produce a standard average normal. We then subtracted the log_2_ ratios of this computed average normal from each tumor. Normalization using GC correction was performed based on the GC content normalization algorithm in HMMcopy (Lai *et al.*, 2016).

## 3 Results

### 3.1 Tangent method overview

We have found systematic biases to be prevalent in both array- and sequencing-based data, both within and across batches, and found that these biases can generate widespread false positive SCNAs that can recur across samples (Figure 1A). In principle, these biases can be overcome by normalizing tumor data only against normal control samples that have been profiled under identical experimental conditions. In practice, many of these experimental conditions are neither known nor measured. We developed the Tangent method to reconstruct normal controls that most accurately represent the tumor noise profile, so as to overcome these tumor-specific biases.

Tangent assumes that variations in experimental conditions can introduce variations in signal intensity or coverage profiles, such that normal samples that represent a single diploid state can produce signal intensity or coverage profiles encompassing a subspace ***N*** of the space of all possible coverage profiles. By accruing a collection of normal samples from the same batch/center as the tumors and with similar noise characteristics, Tangent attempts to construct this reference subspace ***N*** as the space that spans all linear combinations of normal profiles. Tangent then assumes that, for any copy-number profile *T* from a tumor sample, the point in subspace ***N*** that is most similar to *T* represents the profile of a normal sample characterized under similar conditions as *T*. SCNAs are then represented as the difference between *T* and that nearest point in subspace ***N*** (Figures 1B-C; see Methods).

### 3.2 Tangent analysis on microarray data

To assess Tangent’s performance on copy-number profiles generated from microarray data, we applied it to a dataset comprising 497 glioblastomas and 451 normal samples profiled by TCGA using SNP 6.0 arrays. We benchmarked Tangent against two other normalization methods: use of matched normal samples from the same patient (which was possible for only 386 of the GBMs), and use of the five normal samples with noise profiles closest to those in the tumor (Beroukhim *et al.*, 2007). We compared the performance of these normalization approaches in detecting SCNAs based on preservation of signal intensity, reduction in noise, and improvement in signal-to-noise ratio (SNR). We estimated signal as the standard deviation of median signal intensities among all chromosome arms and noise as the median absolute difference between log_2_ copy-ratios of adjacent intervals or probes.

All three normalization methods described preserved signal integrity, but only Tangent normalization consistently reduced systematic noise and thus increased signal-to-noise ratios (Figure 2A-C; Supplementary Figure S2). Normalization using the five nearest normals improves noise levels negligibly, and normalization by matched normals tends actually to increase noise levels and decrease signal-to-noise ratios relative to data that had not been normalized. As a result, segmented copy-number profiles generated after Tangent normalization exhibited less hyper-segmentation than profiles generated using other methods (Supplementary Figure S2).

**Figure 2.**
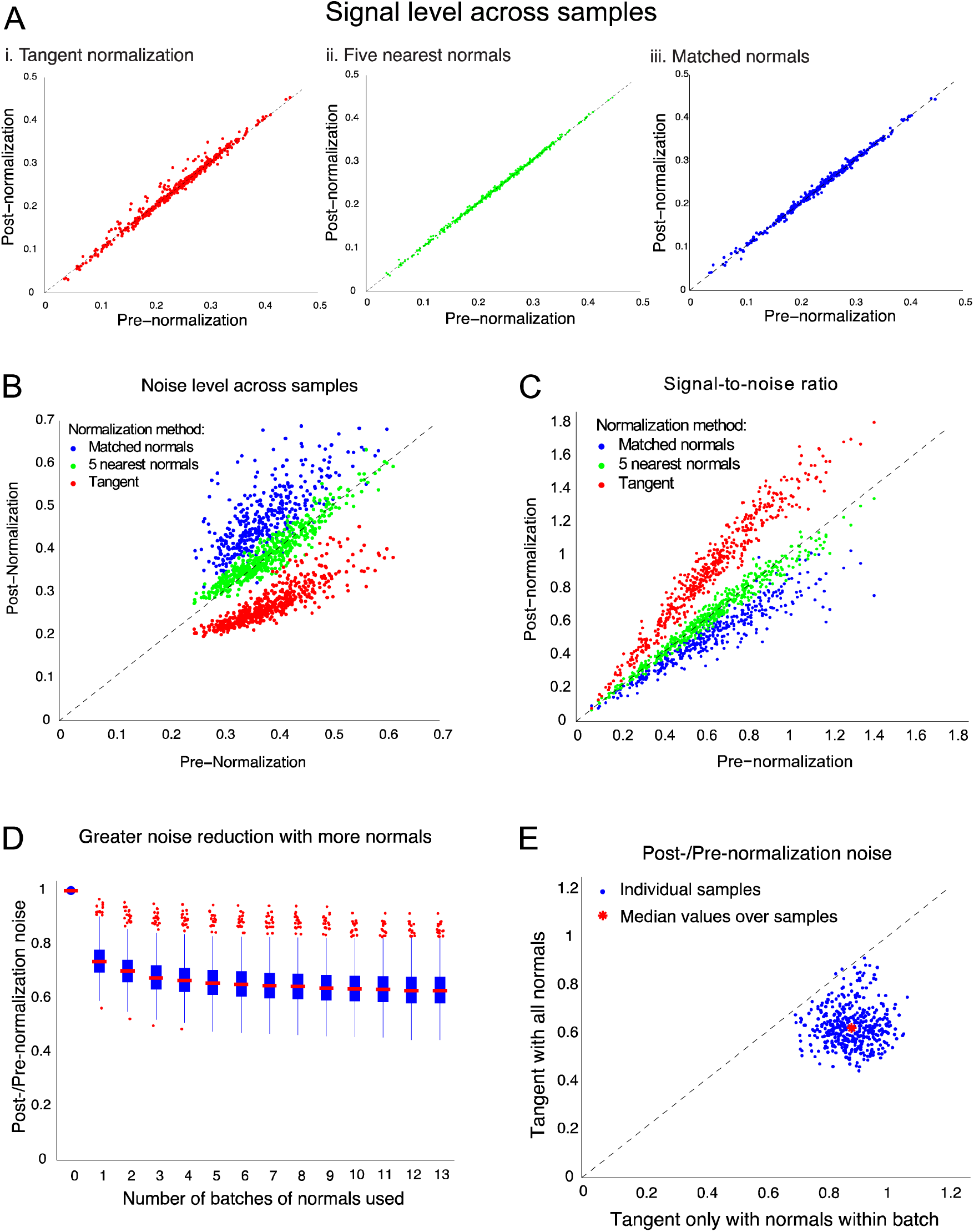
Normalization of SNP array data from 497 TCGA glioblastomas. Scatter plots indicate post-normalization vs. pre-normalization (a) signal, (b) noise level, and (c) signal-to-noise ratios for the normalization methods: Tangent (red), five nearest normals (green), and matched normals (blue). (d) Box plot of post-normalization noise as a fraction of pre-normalization noise, following tangent normalization with increasing numbers of normal samples (approximately 250 normal samples were added in each batch). (e) Noise ratio (post-normalization over pre-normalization noise) for glioblastoma samples following tangent normalization using the entire reference plane vs. tangent normalization using only the normal samples processed in the same batch as a tumor. Almost all samples lie below y=x, indicating that there is greater noise reduction with the full reference plane.

We next investigated the effects of increasing the size of the normal reference pool used by Tangent on reducing noise. We re-applied Tangent to our set of glioblastomas while incrementing the numbers of normal samples used to define our reference subspace from 0 (i.e. no use of Tangent) to 3146 samples across 13 batches (median number of normal samples per batch 255, range 102 to 281). These normal samples represented data generated by TCGA from normal blood leukocytes obtained from patients with a variety of cancers. We observed a monotonic reduction in median noise levels with increasing numbers of normal samples, although this improvement decreased asymptotically and offered negligible benefits after approximately 1000 normal samples (four batches; Figure 2D).

We also investigated the effects of altering the composition of our normal reference pool, and specifically the utility of including normal samples that had been profiled in the same vs. different batches of arrays as the tumor under study. We observed greater noise reduction when utilizing the entire set of normal samples across batches than we did when applying Tangent using a reference subspace containing only normal samples from the same batch (Figure 2E). Nevertheless, whether Tangent utilizes the entire reference subspace or it uses only a subset of normal samples from the same batch, both methods consistently yield lower levels of post-normalization noise compared to pre-normalization noise for all tumors.

### 3.3 Tangent analysis on whole exome sequencing data

We next evaluated Tangent’s performance on sequencing data, by applying it to WES data generated by TCGA from 123 tumors and 129 matched normal samples across four tumor types: low grade gliomas, lung squamous cell carcinomas, prostate adenocarcinomas, and stomach adenocarcinomas (see Supplementary Material). We compared the performance of Tangent with normalizing against matched normals, an average normal from a panel of normals. We also combined each approach with a method that corrects for variations in local GC content (Ha et al., 2014) to determine whether Tangent provides improvements beyond GC correction.

We found that Tangent outperforms these conventional normalization methods. Specifically, the average noise in post-Tangent normalized data is 35% lower than post-normalization against matched normals and 26% lower than post-normalization against an average normal (Figure 3A). This level of noise reduction is attained without significant compromise on signal. The average SNR in post-Tangent normalized data is 58% higher than that post-normalization against matched normals, and 78% higher than post-normalization against an average normal (Figure 3B). Adding GC correction to the other two normalization methods does not enable them to reach the performance of Tangent. Use of Tangent results in 31% lower noise and 57% higher SNR on average than use of matched normals and 55% lower noise and 115% higher SNR on average than use of an average normal. Application of GC-correction to Tangent-normalized data provided only marginal benefit relative to Tangent alone (Figure 3A-B).

**Figure 3.**
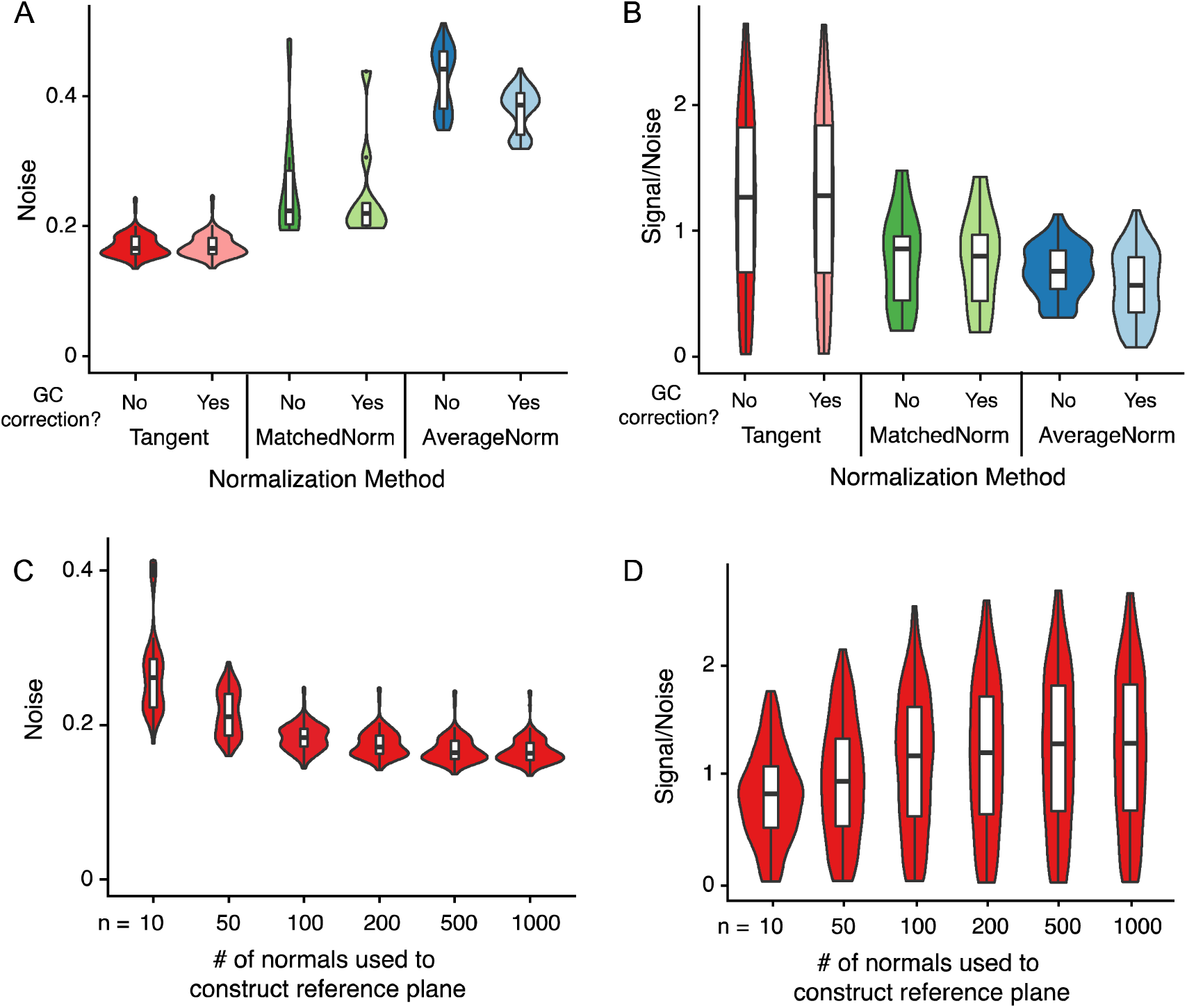
Performance of Tangent on WES data. (a) Noise levels and (b) signal-to-noise ratios for Tangent-normalized data (Tangent); data normalized against corresponding matched normals (MatchedNorm); and data normalized against an average across a panel of normals (AverageNorm), both with and without additional GC correction. (c) Noise and (d) signal-to-noise ratios plotted against the number of normal samples in the reference subspace.

Similar to our experience with SNP arrays, we found that increased numbers of normal samples in the reference pool improved noise profiles after Tangent normalization. We applied Tangent using between 10 and 1000 reference normal samples sequenced by TCGA, including normal samples from patients with the four tumor types under study and six other tumor types (see Supplementary Material). We found that Tangent’s performance plateaued at approximately 200 normal samples (Figure 3C-D).

### 3.4 Pseudo-Tangent: a method to compensate for insufficient normal data

Tangent assumes that systematic noise distributions in tumors are identical to those in normal samples. However, it is often impossible to collect a sufficiently large collection of normal samples to encompass the range of systematic noise types spanned by the tumor samples. In light of this limitation, we developed Pseudo-Tangent as an adaptation of the Tangent pipeline that utilizes a reference subspace composed of signal-subtracted tumor profiles (i.e. “pseudo-normal profiles”) instead of the standard normals used in Tangent. In brief, the method first estimates SCNAs for each tumor using standard Tangent with a limited number of normal samples. Pseudo-Tangent then applies Tangent again to detect SCNAs for each tumor, using a reference subspace comprising other tumors from which the initially detected SCNAs had been subtracted (see Methods).

We applied Pseudo-Tangent to TCGA WES data from 305 Cervical Squamous Cell Carcinoma and Endocervical Adenocarcinoma (CESC) primary tumors. We initially normalized these data against WES data from five normal samples obtained from blood, and used these to generate 305 corresponding pseudo-normal profiles. We then divided the tumors and their matching pseudo-normal profiles into three batches, and normalized each tumor in each batch against the pseudo-normal profiles in the other two batches. (The number of batches is a modifiable parameter.)

We then compared these results to previously generated gold-standard absolute allelic copy-number profiles (Taylor *et al.*, 2018). The gold-standard profiles were generated by applying the standard Tangent pipeline and the ABSOLUTE algorithm (see Methods; Carter *et al.*, 2012) to primarily SNP array data from these 305 tumors and 3,154 normal samples. (ABSOLUTE did, however, use mutation calls from WES data to optimize its tumor purity estimates.) We selected gold-standard profiles based upon a different experimental platform (SNP array data) to minimize cross-contamination of artifacts in the WES data used by Pseudo-Tangent. We measured noise as the average distance of each probe in the Pseudo-Tangent-generated coverage profile from its nearest estimated absolute total copy number level. We found that all 278 CESC tumors displayed lower noise levels after undergoing Pseudo-Tangent normalization than they did after just the initial round of Tangent normalization using only the 5 true normal samples (Figure 4A).

**Figure 4.**
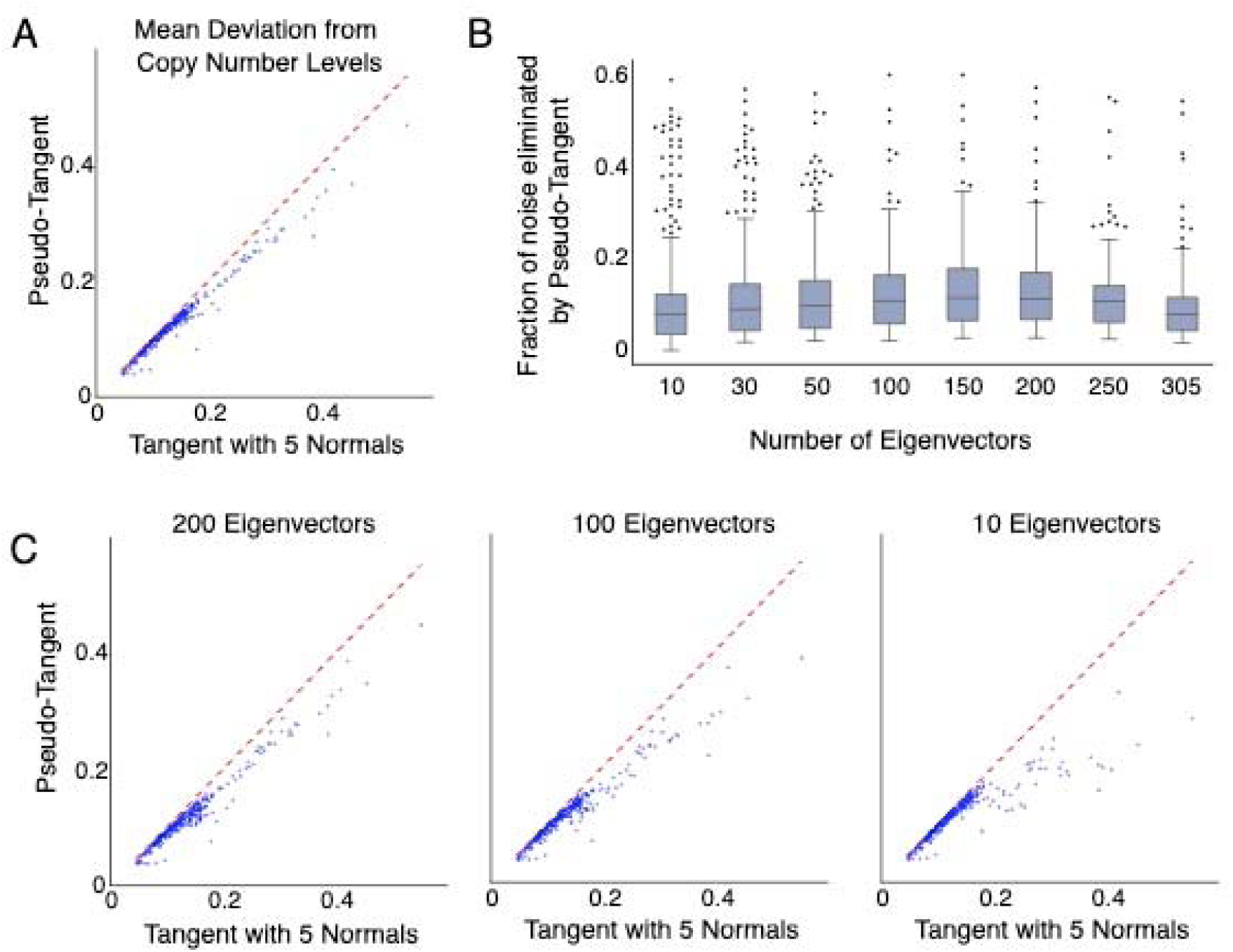
Pseudo-Tangent decreases noise in resulting copy number profiles compared to standard Tangent, as measured by deviation from ABSOLUTE-estimated copy number levels. (a) Average deviation from ABSOLUTE-estimated copy number levels after Pseudo-Tangent (vertical axis) vs. Tangent alone (horizontal axis). (b) Improvement in the deviation from ABSOLUTE-estimated copy number levels after use of Pseudo-Tangent, as a fraction of the deviation after only standard Tangent had been used (vertical axis), against the number of eigenvectors used for Pseudo-Tangent (horizontal axis). The median improvement was greatest when 150 eigenvectors were used. (c) Average deviation from ABSOLUTE-estimated copy number levels after Pseudo-Tangent (vertical axis) vs. Tangent alone (horizontal axis) as in panel (a), after use of (left) 200 eigenvectors, (middle) 100 eigenvectors, and (right) 10 eigenvectors. Although median levels of deviation from ABSOLUTE-estimated copy number levels increased when fewer than 150 eigenvectors were used, the noisiest tumors saw the greatest improvements when only 10 eigenvectors were used.

One concern with applying Pseudo-Tangent is that, with sufficient numbers of pseudo-normal samples, true SCNAs in a tumor may be normalized away due to overfitting. We therefore explored whether limiting the number of dimensions of the pseudo-normal space could improve Pseudo-Tangent’s overall performance. Specifically, we performed eigenvector decomposition of the pseudo-normal reference subspace, retained between 10 to 305 of the eigenvectors with the greatest eigenvalues, and normalized our tumors against a reduced subspace spanned by these eigenvectors. We then determined the number of eigenvectors that provided optimal results, as indicated by generating copy-number profiles with the smallest deviations from the results of our gold-standard ABSOLUTE runs on the same tumors.

We found that the median difference between copy-number levels generated after Pseudo-Tangent and those generated by the gold-standard ABSOLUTE pipeline was lowest when we used the 150 eigenvectors with the greatest eigenvalues (Figure 4B), which captured 98% of the variance of the entire pseudo-normal reference subspace. However, the optimal number of eigenvectors varied across the different tumors. In particular, the noisiest tumors seemed to have greater noise reductions when 10 eigenvectors were used rather than larger numbers of eigenvectors (Figure 4C). This behavior suggests that optimal use of Pseudo-Tangent might take into account the noise level of the tumor being normalized when determining the number of eigenvectors to retain.

## 4 Discussion

Although Tangent was developed for use with SNP array data, we have extended its use to WES data, and in principle it can be applied to any source of copy-number data that measures DNA dosage with varying signal intensity or depth of coverage, such as whole-genome sequencing (WGS) or comparative genomic hybridization (CGH). Indeed, Fehrman et al. developed a similar method to detect SCNAs from transcriptomic profiling data, in which they remove principal components reflecting different transcriptional states to enrich for transcriptional changes reflecting underlying SCNAs (Fehrman *et al.*, 2015). Rearrangements detected by whole genome sequencing (Wala *et al.*, 2018; Rausch *et al.*, 2012; Drier *et al.*, 2013; Layer *et al.*, 2014) can also provide information about copy-number breakpoints, thereby further improving accuracy of SCNA profiles. Further improvements to SNRs can also be obtained from algorithms that determine differences in absolute rather than relative copy-numbers (Carter *et al.*, 2012; Van Loo and Nordgard, 2010). However, these algorithms require normalized copy-number ratios as inputs, and therefore are likely to benefit from the improved normalization Tangent provides.

Accurate SCNA determination relies on having normal control samples that have been processed in identical fashion to the tumors. For example, SCNA profiling of tumors obtained from a large variety of institutional sources--such as may occur when profiling tumors studied in multi-institutional clinical trials or in clinical laboratories--would ideally make use of normal tissue obtained from each institution contributing tumors. Unfortunately, this is often difficult or impossible in practice. Likewise, tumor tissue obtained through careful surgical resection in which the tumor is separated from its blood supply for an extended period may not be adequately reflected by normal DNA from blood samples. Pseudo-Tangent may help remove the effects of systematic noise in these situations by generating pseudo-normals from tumors that were processed in similar fashion to each other. However, application of Pseudo-Tangent carries risk of overfitting and loss of signal, particularly if SCNAs are not adequately removed while generating pseudo-normals from tumor samples. In situations where true normals are available, extensive profiling of these normals as controls for the tumors is preferable to computational generation of pseudo-normals as described here.

The Tangent pipeline we describe here was the basis for copy-number determination across TCGA. Additionally, both Tangent and Pseudo-Tangent are widely applicable to a large variety of research and clinical applications and copy-number profiling platforms, and can be integrated with further improvements in SCNA detection that make use of alternative sources of information such as rearrangement locations and tumor purity and ploidy.

## 5 Acknowledgements

We are grateful for support from our Broad Institute colleagues in the Genomics Platform and colleagues from The Cancer Genome Atlas Project. Stefano Monti, Jeffrey Gentry, Bryan C. Hernandez, Michael O’Kelly, Marc-Danie Nazaire, Nam H. Pho, Travis I. Zack, Nicholas Stransky, Joshua Gould, David Twomey, Mark Nadel, Wendy Winckler, contributed helpful discussions and analysis support.

