## Supplementary Figures and Methods for "The Tangent copy-number inference pipeline for cancer genome analyses"

### **8 Supplementary Figures**

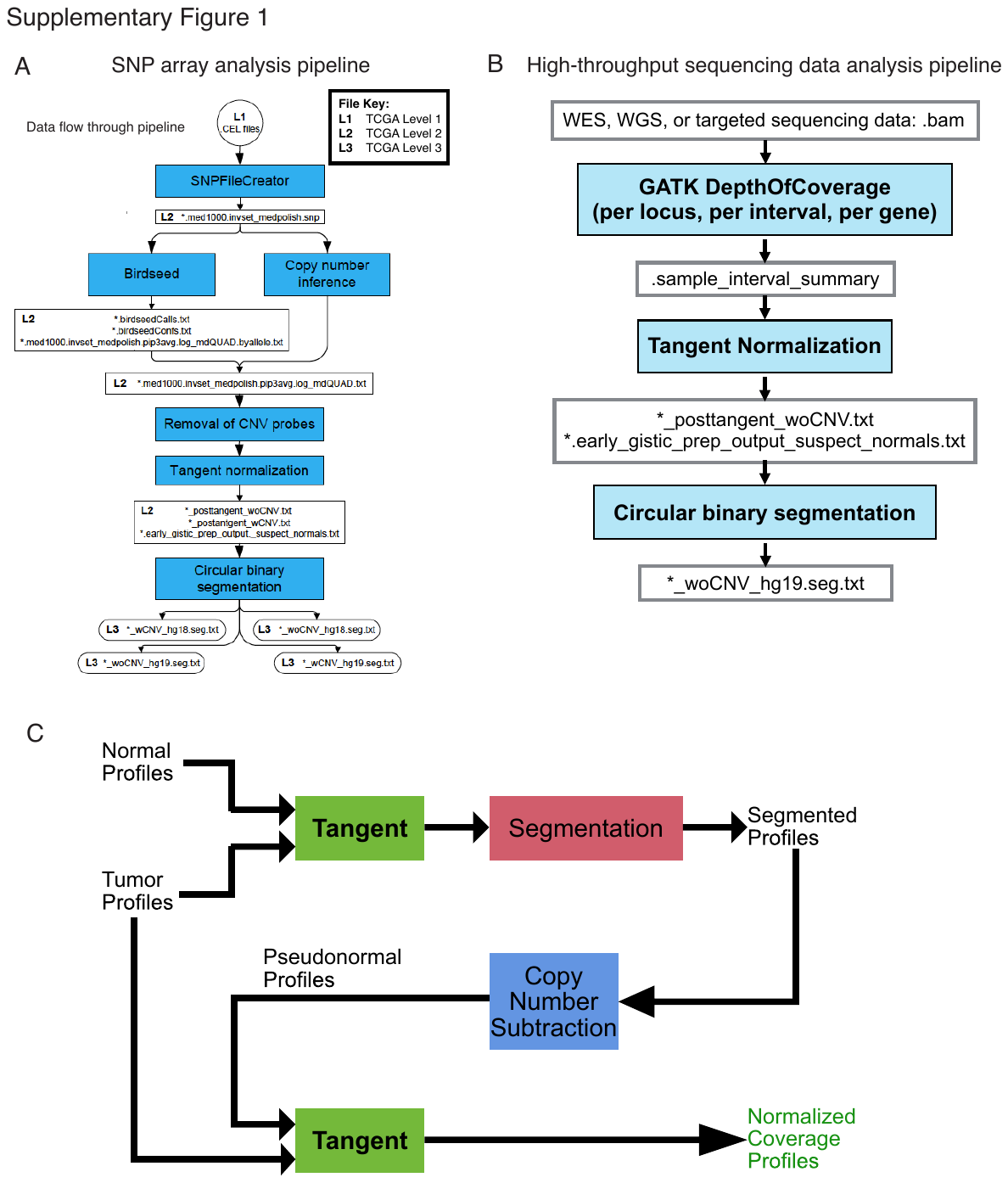
**Supplementary Figure S1.** (a) Overview of the Tangent copy number inference pipeline for SNP array data. Schematic view of pipeline with key output files for pipeline modules and assigned TCGA levels included. (b) Overview of Tangent copy number inference pipeline for high-throughput sequencing data. (c) Overview of the Pseudo-Tangent pipeline.

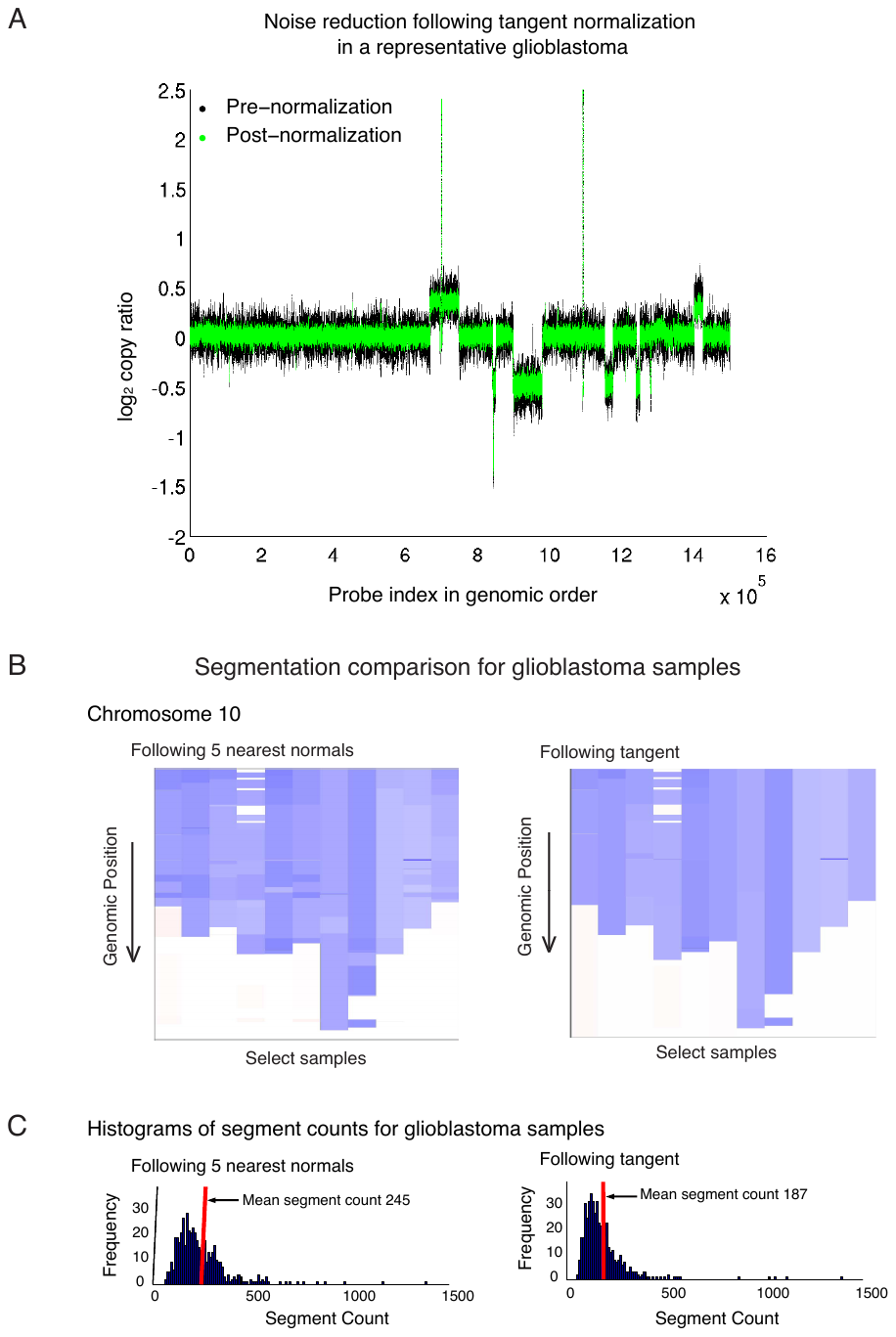
**Supplementary Figure S2.** (a) Noise reduction following tangent normalization in a representative glioblastoma tumor. 100-marker moving average of log_2_ copy-ratios are shown for a representative glioblastoma sample across the autosomes before (black) and after (green) tangent normalization. A moving average is employed for visualization purposes only. (b) Post-segmentation results for selected glioblastoma samples following (left) 5NN and (right) Tangent normalization. White is copy-neutral. Blue indicates a deletion with intensity of color increasing as the copy-number decreases. Samples are displayed in the same order in both panels. Less hypersegmentation is observed when CBS is applied to tangent-normalized data. (c) Histograms of segment counts for 497 TCGA glioblastoma tumors when CBS follows (left) 5NN and (right) tangent. Decreased segments counts for tangent normalized data is consistent with decreased hypersegmentation.

### **9 Supplementary Data**

#### **9.1 Preparation of SNP array data**

Our pipeline was the basis for analyses of all Affymetrix SNP 6.0 array data for The Cancer Genome Atlas (TCGA). The TCGA .CEL file inputs and all pipeline outputs are available at the GDC <https://gdc-portal.nci.nih.gov/legacy-archive> as Level 1 Data. Output files of modules (1) – (4) are available as Level 2 Data; normalized, segmented data (outputs of module (5)) are uploaded as Level 3 Data (Supplementary Figure 1). We construct the standard reference plane using all TCGA normal samples obtained from blood that pass Levels 1 and 2 of quality control.

Affymetrix SNP 6.0 arrays contain 906,600 “SNP markers” associated with single nucleotide polymorphisms and 946,000 “copy number markers” at other locations (<http://tools.thermofisher.com/content/sfs/brochures/cn_snp_variation_technote.pdf>). Each genomic locus and SNP allele is represented by multiple probes on these arrays (a “probeset”). Probe-level intensities are represented in .CEL files generated by Affymetrix GeneChip Command Console. We use SNPFileCreator, a Java implementation of the dChip signal intensity determination algorithm, to normalize and merge intensity values for each probeset (Li and Wong, 2001b; 2001a). Probe intensities across a sample are first scaled to achieve a median brightness value of 1000 and then are subjected to quantile normalization. The normalized probe intensities of each sample are mapped to a reference sample using model-based expression indices. Median polish is then applied to each probeset across samples to produce one value per probeset for each sample. We apply SNPFileCreator to all arrays in a batch, as defined by the joint PCR amplification step.

#### **9.2 Calibration of CN Markers and CN inference**

Probeset intensities are mapped to copy-number levels (“calibrated”) on a batch-by-batch basis, assuming a linear relationship between signal intensity and copy-number. SNV probes are calibrated based on A and B intensities observed within the current and prior batches, while the CNV probes are calibrated based on an experiment using cell lines.

Calibration of a probeset is determined by two parameters: the background signal intensity and a scale factor that specifies the change in intensity resulting from each added copy of DNA. For SNP loci, Birdseed is used to calibrate probesets for each allele, using intensity data collected from normal samples, allele-specific background, and scale parameters (Korn et al., 2008). The resulting copy-numbers for the two alleles are summed to obtain total copy-number estimates.

Our calibration of the CN markers relies on a two-step modeling approach (“copy-number inference”) based on SNP6.0 array data of an X-dosage experiment performed on 46 samples from 5 cell lines with known variation of copy number of the X-chromosome from 1 to 5. We first calibrate each probeset on X by applying linear regression to the experimental data to fit the parameters $\beta_{i0}$ and $\beta_{i1}$ of the model :

**(i)** $\boldsymbol{I}_{\boldsymbol{i}}\boldsymbol{=}\boldsymbol{\beta}_{\boldsymbol{i0}}\boldsymbol{+}\boldsymbol{\beta}_{\boldsymbol{i1}}\boldsymbol{C}_{\boldsymbol{i}}$**.**

Here variable $\boldsymbol{I}_{\boldsymbol{i}}$ represents the intensity of probeset i, variable $\boldsymbol{C}_{\boldsymbol{i}}$ represents the copy level at probeset i, and parameters $\beta_{\boldsymbol{i0}}$ and $\beta_{\boldsymbol{i1}}$correspond respectively to the background signal intensity and the scale factor that specifies the change in intensity resulting from each added copy of DNA.

We extend the resulting calibration of the X probesets to a calibration for all probesets across the genome by modeling the background signal intensity and the scale factor as functions of local sequence features and median intensity across samples as follows:

**(ii)** $\beta_{\boldsymbol{ik}}\boldsymbol{=}\boldsymbol{\alpha}_{\boldsymbol{k0}}\boldsymbol{+}\boldsymbol{\alpha}_{\boldsymbol{k1}}\boldsymbol{G}\boldsymbol{C}_{\boldsymbol{i}}\boldsymbol{+}\boldsymbol{\alpha}_{\boldsymbol{k2}}\boldsymbol{F}{\boldsymbol{L}^{\boldsymbol{(sty)}}}_{\boldsymbol{i}}\boldsymbol{+}\boldsymbol{\alpha}_{\boldsymbol{k3}}\boldsymbol{F}{\boldsymbol{L}^{\boldsymbol{(nsp)}}}_{\boldsymbol{i}}\boldsymbol{+}\boldsymbol{\alpha}_{\boldsymbol{k4}}{\boldsymbol{I}_{\boldsymbol{i}}}^{\boldsymbol{m}}\boldsymbol{+}\boldsymbol{\alpha}_{\boldsymbol{k5}}{\boldsymbol{(I}_{\boldsymbol{i}}}^{\boldsymbol{m}}\boldsymbol{)}^{\boldsymbol{2}}$

for $k \in\{0,1\}$. The variable $\boldsymbol{G}\boldsymbol{C}_{\boldsymbol{i}}$ represents the GC content of the i^th^ probeset, $\boldsymbol{F}{\boldsymbol{L}^{\boldsymbol{(sty)}}}_{\boldsymbol{i}}$and $\boldsymbol{F}{\boldsymbol{L}^{\boldsymbol{(nsp)}}}_{\boldsymbol{i}}$ represent the fragment lengths of the STY and NSP fragments of the i^th^ probeset respectively, and ${\boldsymbol{I}_{\boldsymbol{i}}}^{\boldsymbol{m}}$ represents the probeset median intensity across the samples. The linear dependence on GC content and fragment lengths and quadratic dependence on median intensity provides a good fit to our X-dosage array data. We apply linear regression again, this time to find the parameters $\alpha_{\boldsymbol{kl}}$for $k\in\{0,1\}$ and $l\in\{0,1,2,3,4,5\}$ that best fit model (ii). The parameters $\alpha_{\boldsymbol{kl}}$ are independent of the probeset and, for k = 1 or 2, this regression is performed collectively on the complete set of X-chromosome data, $\{\beta_{ik}, GC_{i}, F{L^{(sty)}}_{i}, F{L^{(nsp)}}_{i}, {I_{i}}^{m}{\}}_{i\in I_{x}}$ where $I_{x}$ is the collection of indices for the X probesets. Each of the parameters for both models (i) and (ii) described above is computed once based on the results of the X-dosage experiment. Model (ii) is then used to predict the background and scale factor across the genome for each new batch of SNP6.0 array data. While GC content and fragment lengths do not vary with the batch, the median intensity must be computed separately for each batch.

#### **9.3 Datasets used in Tangent WES analysis**

The datasets used in Tangent WES analysis contained 123 tumors and 129 matched normal samples (123 normal blood and 6 normal tissue samples) across four cancer types:

- STAD (Stomach Adenocarcinoma): 57N 51T
- LUSC (Lung Squamous Cell Carcinoma): 48N 48T
- LGG (Lower Grade Glioma): 14N 14T
- PRAD (Prostate Adenocarcinoma): 10N, 10T

We also constructed reference planes using 10-1000 non-matched normals from 10 different cancer types in TCGA: CESC, GBM, KIRC, LAML, LGG, LUSC, PRAD, READ, STAD, UCEC.
